# C9orf72 Dipeptide Repeats Inhibit UPF1-Mediated RNA Decay Independent of Stress Granule Formation

**DOI:** 10.1101/623769

**Authors:** Yu Sun, Aziz Eshov, Junjie U. Guo

**Affiliations:** Department of Neuroscience, Yale University School of Medicine, New Haven, CT 06520, USA; Program in Cellular Neuroscience, Neurodegeneration and Repair, Yale University School of Medicine, New Haven, CT 06520

## Abstract

Expansion of an intronic (GGGGCC)_n_ repeat region within the *C9orf72* gene is a major cause of familial amyotrophic lateral sclerosis and frontotemporal dementia (c9ALS/FTD). A pathological hallmark in c9ALS/FTD is the accumulation of misprocessed RNAs, which are often targets of RNA surveillance pathways in normal cells. Here we show that nonsense-mediated decay (NMD) and other RNA decay mechanisms involving upstream frameshift 1 (UPF1), collectively referred to as UPF1-mediated RNA decay (UMD), are broadly inhibited in c9ALS/FTD brains. These effects are recapitulated in cultured cells by the ectopic expression of arginine-rich dipeptide repeats (DPRs), poly(GR) and poly(PR). Despite these two DPRs causing the recruitment of UPF1 to stress granules, stress granule formation is neither sufficient nor necessary for UMD inhibition. Our results suggest that UMD inhibition may accelerate the accumulation of deleterious RNAs and polypeptides in c9ALS/FTD.

## INTRODUCTION

Since the identification of a GGGGCC (G_4_C_2_) repeat expansion within the first intron of *C9orf72* as the major cause of both familial ALS and FTD^1,2^, a variety of mechanisms, including haploinsufficiency^3^, RNA toxicity^4^, and dipeptide repeat (DPR) toxicity^5–7^, have been proposed to explain the pathogenicity of this autosomal dominant mutation. The expanded G_4_C_2_ repeat region is transcribed in both directions, producing the C9orf72 pre-mRNA, which contains intronic G_4_C_2_ repeats, and the antisense RNA that contains G_2_C_4_ repeats^8^. Both RNAs accumulate in nuclear foci and, once exported, can be translated into distinct sets of DPR-containing polypeptides^8^.

In addition to *C9orf72*, genes encoding RNA-binding proteins such as FUS (fused in sarcoma) and TDP-43 (TAR DNA-binding protein 43) are also enriched in ALS/FTD-associated mutations, suggesting that RNA misprocessing may be a converging point in pathophysiology^9,10^. Indeed, widespread RNA processing defects have been described by previous high-throughput RNA sequencing (RNA-seq) studies using post-mortem brain tissues from both sporadic ALS (sALS) and c9ALS/FTD patients^4,11^. In normal cells, misprocessed mRNAs are targeted for degradation by multiple RNA surveillance pathways^12–14^, such as nonsense-mediated decay (NMD), no-go decay (NGD), and nonstop-mediated decay (NSD). The accumulation of aberrant RNAs in c9ALS/FTD brains hints at the possibility that one or more mRNA surveillance pathways may be compromised.

## RESULTS

### UPF1-mediated RNA decay targets accumulate in c9ALS brains

We began by re-examining the post-mortem c9ALS, sALS, and control RNA-seq data^11^ from the frontal cortex, which shows both RNA foci and DPR pathology in c9ALS/FTD. As shown previously, intron retention events were prevalent among c9ALS subjects, as indicated by a right-shift in the cumulative distribution function (CDF) of the fraction of reads mapped to the introns of each gene (Figure 1A, red curve), but not in sALS subjects (Figure 1A, blue curve). Intron-retaining mRNAs typically contain premature stop codons, which would render them substrates for NMD. To assess whether NMD may be inactivated in c9ALS/FTD, we next quantified the expression of endogenous regulatory targets of NMD. When we compared the abundance of previously defined (*N*=76) endogenous NMD targets^15^ between ALS and controls, we found that they were broadly up-regulated in c9ALS (Figure 1B, compare red and black curves), but not sALS subjects (Figure 1B, compare blue and black curves). These observed changes were replicated when we used an expanded list (*N*=1,271) of NMD targets^16^ (Figure 1B, orange and light blue curves). In addition, we found that the mRNAs for key NMD factors, UPF3B, SMG5, and SMG6 were significantly up-regulated in c9ALS subjects (Figure 1C), consistent with the auto-regulatory feedback upon NMD inhibition^17^.

**Figure 1.**
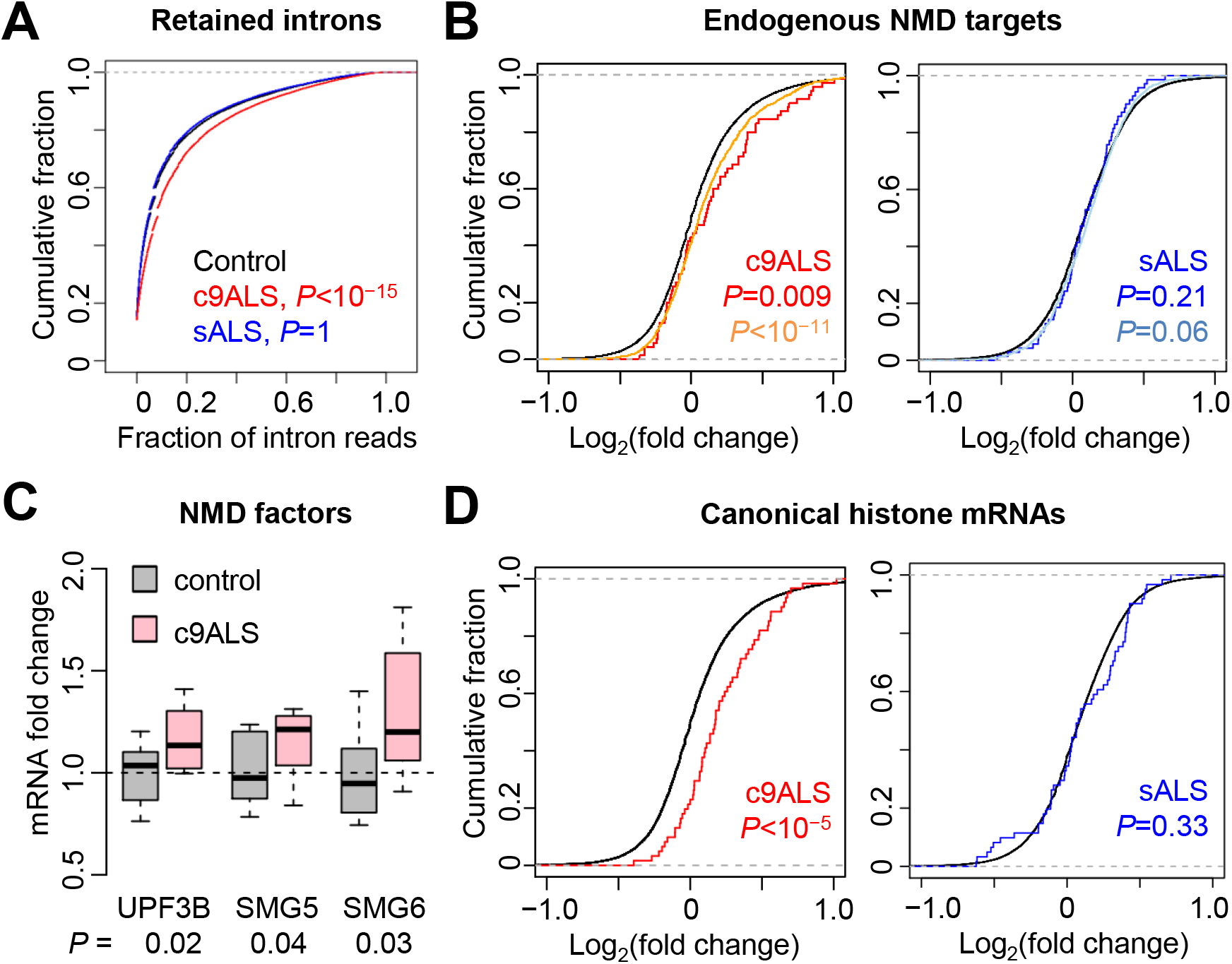
UMD targets accumulate in c9ALS brains. (**A**) CDFs of the fraction of intron-mapping reads for each gene. (**B**) CDFs of changes in RNA abundance for all genes (black), NMD targets identified in Tani et al. (*N*=76, red and blue), and NMD targets identified in Colombo et al. (*N*=1,271, orange and light blue), comparing between c9ALS and controls (*left*) or sALS and controls (*right*). (**C**) Changes in mRNA abundance of known NMD factors between c9ALS and controls. Expression levels were normalized to the means of control subjects. *P* values, one-sided Student’s t-tests. (**D**) CDFs of changes in RNA abundance for all genes (black) and canonical histone genes (*N*=87, red and blue), comparing between c9ALS and controls (*left*) or sALS and controls (*right*). All *P* values were from one-sided Mann-Whitney-Wilcoxon tests except in Figure 1C.

Canonical histone mRNAs lack polyA tails and instead have 3ʹ end stem-loop structures^18^. Histone mRNA decay requires UPF1, but not other NMD factors^18,19^. Similar to NMD targets, we observed an overall up-regulation of canonical histone mRNAs in c9ALS, but not in sALS subjects (Figure 1D). In contrast, noncanonical, polyadenylated histone variant mRNAs were largely unchanged between c9ALS and controls (Figure 1–figure supplement 1), suggesting that accumulation is specific to those UPF1-dependent, canonical histone mRNAs.

Collectively, the accumulation of intron-retaining mRNAs, NMD regulatory targets, NMD factor mRNAs, and canonical histone mRNAs suggest that multiple UPF1-mediated RNA decay mechanisms, hereafter collectively referred to as UMD, are broadly inactivated in c9ALS/FTD.

### Arginine-rich DPRs acutely inhibit UMD in cultured cells

The *C9orf72* repeat expansion can be transcribed in both directions, producing both sense G_4_C_2_ repeat- and antisense G_2_C_4_ repeat-containing RNAs^8^, which can be translated into a total of five DPRs: poly(GA), poly(GP), poly(GR), poly(PA), and poly(PR). To test whether any of these gene products may be sufficient to inhibit UMD, we ectopically expressed each of the 2 repeat RNAs as part of the 5ʹ untranslated regions (UTRs) of a GFP transcript, and each of the 4 codon-optimized DPRs as GFP fusion proteins^20^ in HEK293 cells, and quantified the expression levels of several UMD targets that accumulated in c9ALS subjects, including three NMD targets, two NMD factor mRNAs, and a canonical histone mRNA. We failed to validate the codon-optimized poly(GP) expression construct, and excluded it from our analysis. Interestingly, poly(GR) and poly(PR) increased the expression levels of six and five out of six tested UMD targets, respectively, within 24 hours after transfection (Figure 2A), whereas no significant changes in UMD target expression were detected in cells expressing repeat RNAs nor the two alanine-rich DPRs (Figure 2A),

**Figure 2.**
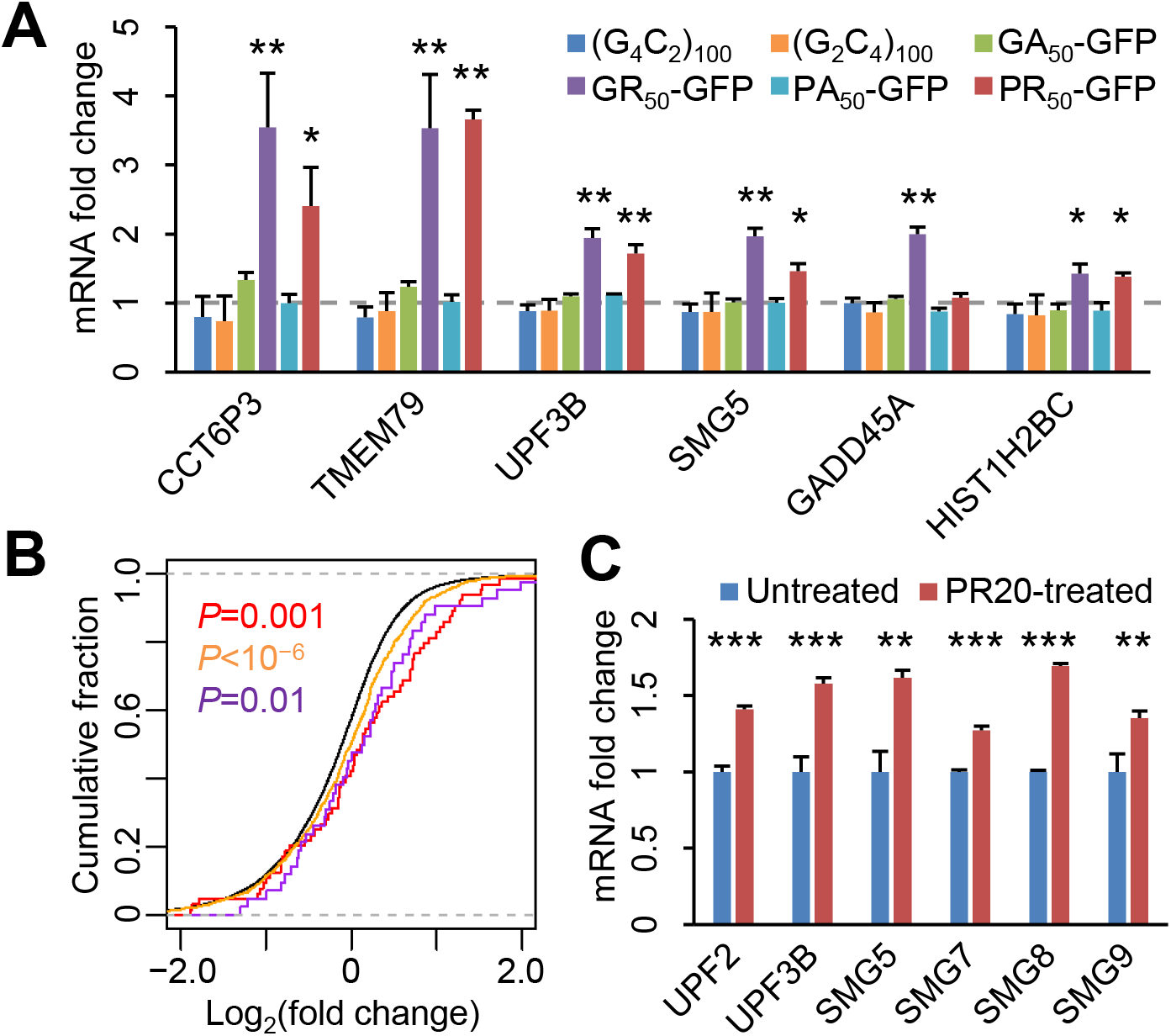
Arginine-rich DPRs acutely inhibit UMD in cultured cells. (**A**) Changes in mRNA abundance of six UMD targets in HEK293 cells ectopically expressing each of the two repeat RNAs and four DPRs. Expression levels relative to GAPDH were normalized to the GFP-only control. Error bars, standard deviations. *, *P*<0.05; **, *P*<0.01, one-way ANOVA with post-hoc Tukey HSD test. (**B**) CDFs of changes in RNA abundance for all genes (black), NMD targets identified in Tani et al. (red), NMD targets identified in Colombo et al. (orange), and canonical histone mRNAs (purple), comparing between PR_20_-treated and control K562 cells. *P* values, one-sided Mann-Whitney-Wilcoxon tests. (**C**) Changes in mRNA abundance of NMD factors between control (*N*=3) and PR_20_-treated (*N*=3) K562 cells. Error bars, standard deviations. **, *P*<0.01; ***, *P*<0.001, one-sided Student’s t-tests.

To validate this finding on a transcriptome scale, we examined the previously published RNA-seq data from K562 leukemia cells treated with synthetic PR_20_ peptides^21^. Again, we observed a substantial upregulation of both NMD targets (Figure 2B, red and orange curves) and canonical histone mRNAs (Figure 2B, purple curves). Multiple NMD factors were up-regulated (Figure 2C). Collectively, these results suggest that poly(GR) and poly(PR) are sufficient to cause acute UMD inhibition in cultured cells.

### UMD inhibition is independent of stress granule recruitment of UPF1

Previous studies have linked the cellular toxicity of poly(GR) and poly(PR) to their influence on stress granule dynamics^22–24^. To test whether UMD inhibition may be linked to stress granule formation, we examined the localization of UPF1 and G3BP1, a stress granule marker, in HeLa cells expressing each repeat RNA or DPR. In control cells, UPF1 exhibited diffuse cytoplasmic localization, with G3BP1-negative small foci that resembled P-bodies (Figure 3A). In contrast to previous studies^25,26^, we did not observe a significant increase in stress granules in cells that expressed either G_4_C_2_ or G_2_C_4_ repeat RNAs (Figure 3A & B), suggesting that stress granule formation may not be a direct consequence of repeat RNA expression. Poly(GA) and poly(PA) expression did not increase stress granules, either. On the contrary, poly(GR) and poly(PR) induced the concentration of UPF1 in G3BP1-positive stress granules in substantial fractions (25-50%) of cells (Figure 3A & B). These results are consistent with previous studies on the influence of poly(GR) and poly(PR) on stress granule assembly^22–24^ and the presence of UPF1 in stress granules^27^, raising the possibility that UMD inhibition may be due to the sequestration of UPF1 and other UMD factors in stress granules.

**Figure 3.**
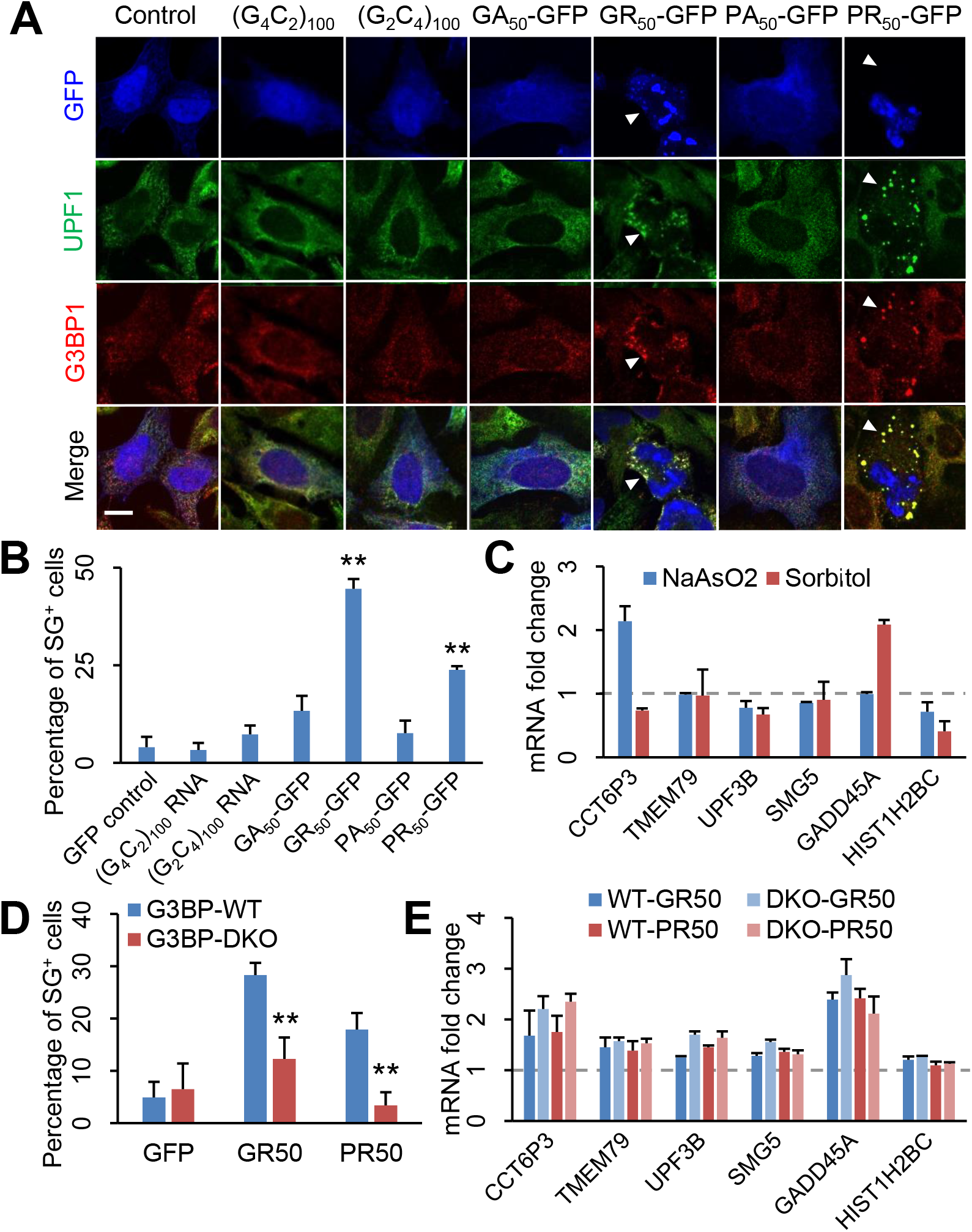
UMD inhibition is independent of stress granule recruitment of UPF1. (**A**) UPF1 and G3BP1 localization in HeLa cells expressing each of the two repeat RNAs and four DPRs. Stress granules are indicated by arrowheads. Scale bar, 10 μm. (**B**) Quantification of stress granules in repeat RNA- or DPR-expressing cells. **, *P*<0.01, one-way ANOVA with post-hoc Tukey HSD test. (**C**) Changes in mRNA abundance of UMD targets in U2 O-S cells treated with 0.4M sorbitol or 0.2mM NaAsO_2_. Expression levels relative to GAPDH were normalized to the untreated control. (**D**) Quantification of stress granule formation in G3BP-WT and -DKO cells. **, *P*<0.01, Student’s t-tests. (**E**) Changes in UMD target RNA abundance in G3BP-WT and -DKO cells. Expression levels relative to GAPDH were normalized to the GFP-only control. All error bars represent standard deviations.

To test whether DPRs can also induce stress granules in neurons, we expressed each DPR in mouse primary cortical neurons. Compared to those in cell lines, Upf1-positive stress granules were exceedingly rare in neurons (Figure 3–figure supplement 1). We detected Upf1-positive, Ataxin-2-positive stress granules only in a small fraction (<5%) of poly(GR)-expressing neurons (Figure 3–figure supplement 1), but not in neurons expressing other DPRs, consistent with a recent study showing that poly(GR) co-localizes with stress granule-like inclusions in (G_4_C_2_)_149_-expressing mouse brain^28^. These results suggest either that the neuronal cytoplasm may be less permissive to stress granule formation, or that neuronal stress granules may exhibit distinct morphology and/or dynamics from those in cell lines.

To directly test the impact of stress granule assembly on UMD, we induced stress granules by applying oxidative stress or osmotic stress using sodium arsenite (NaAsO_2_) or sorbitol, respectively. We confirmed that UPF1 and, to a lesser extent, UPF3B, were both recruited to arsenite-induced stress granules (Figure 3 –figure supplement 2). Even though arsenite and sorbitol induced stress granules in nearly all cells (Figure 3 –figure supplement 2), arsenite caused the upregulation of only one out of six UMD targets (Figure 3C), whereas sorbitol upregulated a different target, suggesting that neither arsenite-nor sorbitol-induced stress granules are sufficient to broadly inhibit UMD in cells.

To test whether stress granule formation may be required for UMD inhibition, we ectopically expressed GFP, poly(GR), or poly(PR) in wild-type (G3BP-WT) or G3BP1/2 double-knockout (G3BP-DKO) U-2 OS cells^29^. Consistent with previous studies^22,23^, poly(PR)-induced stress granule formation was abolished in G3BP-DKO cells (Figure 3D). Interestingly, in G3BP-DKO cells, poly(GR) promoted the formation of UPF1-positive, TIAR-positive stress granules that were fewer and smaller in size compared to those in G3BP-WT cells (Figure 3D; Figure 3–figure supplement 3). Despite the large reduction and morphological changes of stress granules in G3BP-DKO cells, UMD target upregulation by poly(GR) and poly(PR) was largely unchanged and in some cases (e.g. UPF3B) even slightly increased (Figure 3E). These results indicate that stress granule assembly is neither sufficient nor required for DPR-induced UMD inhibition, rejecting the notion that stress granule assembly may contribute to UMD inhibition^30^.

## DISCUSSION

Emerging evidence suggests that widespread transcriptomic aberration is a distinguishing feature of c9ALS/FTD. Previous studies have largely attributed it to the deficiency of splicing factors (e.g. hnRNP H), leading to increased production rates of aberrant RNAs^4,11^. Our results suggest that decreased degradation rates resulting from NMD inhibition also contribute to the accumulation of potentially deleterious RNAs in c9ALS/FTD.

Aside from targeting misprocessed RNAs such as intron-retaining mRNAs, a separate function of NMD is lowering the abundance of many endogenous, properly processed RNAs. We found in c9ALS subjects an overall upregulation of numerous NMD target mRNAs as well as many noncoding RNAs including pseudogene transcripts and the splice variants of small nucleolar RNA precursors^31^ (data not shown). Since many NMD factors including UPF1 are conserved in all eukaryotes and required for survival in most metazoans^32–34^, chronic NMD inhibition would likely be detrimental to neuronal viability^13^.

Our analysis shows that inhibition of RNA decay is not limited to NMD, but also affects other UPF1-dependent decay substrates such as canonical histone mRNAs^18,19^. Notably, aberrations in heterochromatin have been recently linked to poly(PR) expression in mouse models^35^. It remains to be determined whether the accumulated histone mRNAs are translated, and how they may impact chromatin modifications and neuronal survival in c9ALS/FTD.

Ectopically expressed arginine-rich DPRs are sufficient to recapitulate UMD inhibition in cultured cells, suggesting that in c9ALS/FTD patients UMD inhibition may result from low levels of poly(GR) and/or poly(PR) accumulation over a long period. Considering that arginine-rich DPRs are strong stress granule inducers and that UPF1 is a known stress granule component, it is tempting to hypothesize that these DPRs inhibit UMD by sequestering UPF1 in stress granules and away from P-bodies and the rest of cytoplasm, where UMD can occur efficiently. However, we present multiple lines of evidence arguing against this hypothesis: (i) In cell lines, poly(GR) promoted stress granule assembly more strongly than poly(PR), but these DPRs inhibited UMD to similar degrees. (ii) DPR expression rarely caused stress granule assembly in neurons. (iii) Although chemically induced stress granules also recruited UPF1, they did not broadly inhibit UMD. (iv) Blocking stress granule assembly by deleting G3BP1/2 did not alleviate UMD inhibition. These results collectively reject the notion that stress granule assembly contributes to UMD inhibition. We speculate that UMD inhibition may result from the inhibitory effect of arginine-rich DPRs on global translation^36,37^. Without translation, NMD targets would not be distinguishable from non-targets. It remains to be tested whether the inhibition of other UMD mechanisms may result secondarily from translational arrest-induced NMD inhibition.

A recent study has reported that ectopic expression of NMD factors can alleviate poly(GR)- and poly(PR)-induced toxicity in SH-SY5Y neuroblastoma cell line and in flies^30^. In that study, overexpression of UPF1 shows stronger suppression of toxicity than that of UPF2, consistent with our finding that RNA decay pathways other than NMD are also involved. Considering that UPF1 and UMD have now been implicated in multiple ALS/FTD models^30,38,39^, restoring UMD activity to a normal level may represent a promising therapeutic approach.

## MATERIALS AND METHODS

### RNA-seq data analysis

FASTQ files of previously published RNA-seq datasets were downloaded from European Nucleotide Archive (ENA), and uniquely mapped to the human (GRCh38) or mouse (GRCm38) reference genome using STAR. Exon/intron reads for all annotated genes were quantified using BEDTools. After excluding genes with <1 read/sample on average, reads per million uniquely mapped reads (RPM) values were calculated for each gene. A pseudo-RPM value of 0.1 was added to all RPM values before calculating the fold change between ALS or experimental samples and controls. For NMD target analysis, two independent gene sets were used: (1) “group C” genes in Tani et al. (*N*=76), (2) genes with *P*_meta_meta_<10^−5^ in Colombo et al. (*N*=1,271). Expression changes of each gene set were compared to all genes using one-sided Mann-Whitney-Wilcoxon tests.

### RT-qPCR analysis

Expression constructs for repeat RNAs or GFP-tagged codon-optimized DPRs were transfected in HEK293 cells or U-2 OS cells with Lipofectamine 2000 (Invitrogen) or FuGENE HD (Promega), respectively, according to the manufacturers’ instructions. 24−48 hrs after transfection, total RNA was extracted with TRIzol and treated with Turbo DNase (Invitrogen). RT-qPCR was performed on a CFX96 RT-PCR system (Bio-Rad) with Luna Universal One-Step RT-qPCR reagents (New England Biolabs). 50 ng total RNA and 250 nM primers were added in each reaction. Ct values were averaged across 2 technical replicates and normalized to GAPDH internal controls. 3 biological replicates were performed.

### Immunocytochemistry

Transfected HEK293 cells, HeLa cells, U-2 OS cells, or E18 mouse cortical neurons on coverslips were fixed at room temperature with 4% paraformaldehyde in PBS, washed three times with PBS, and permeabilized with 0.5% Triton X-100. After washing and 1 hr blocking with 10% goat serum, cells were incubated with primary antibodies (rabbit anti-UPF1, Cell Signaling Technology #12040; mouse anti-G3BP1, Millipore #05-1938; rabbit anti-TIAR, BD Biosciences #610352; rabbit anti-Ataxin-2, BD Biosciences #611378) diluted in 5% BSA at 4°C overnight. After washing, cells were incubated for 1 hr at room temperature with Alexa Fluor-conjugated secondary antibodies (Invitrogen) diluted in 5% BSA. After washing, cells were stained with Hoechst 33342, mounted with ProLong Diamond Antifade Reagent, and imaged on a Nikon Ti-E Eclipse inverted microscope (spinning disc confocal) with a 60X oil objective.

## ACKNOWLEDGEMENTS

We thank D. Trotti for DPR expression constructs, N. Kedersha and P. Anderson for G3BP-WT and G3BP-DKO cell lines, C. Reddy and P. Gopal for assistance in primary neuronal culture, J. Steitz, A. Alexandrov, A. Horwich, J. Lim, S. Chandra, and members of the Guo lab for discussions and comments on the manuscript. This work was supported by an NIH New Innovator Award (DP2GM132930-01), the Muscular Dystrophy Association (MDA602934), the Ludwig Family Foundation, and the Yale Scholar in Neuroscience Fund. J.U.G. is a NARSAD Young Investigator and a Klingenstein’simons Fellow in Neuroscience.

---

**Figure 1 – figure supplement 1.**
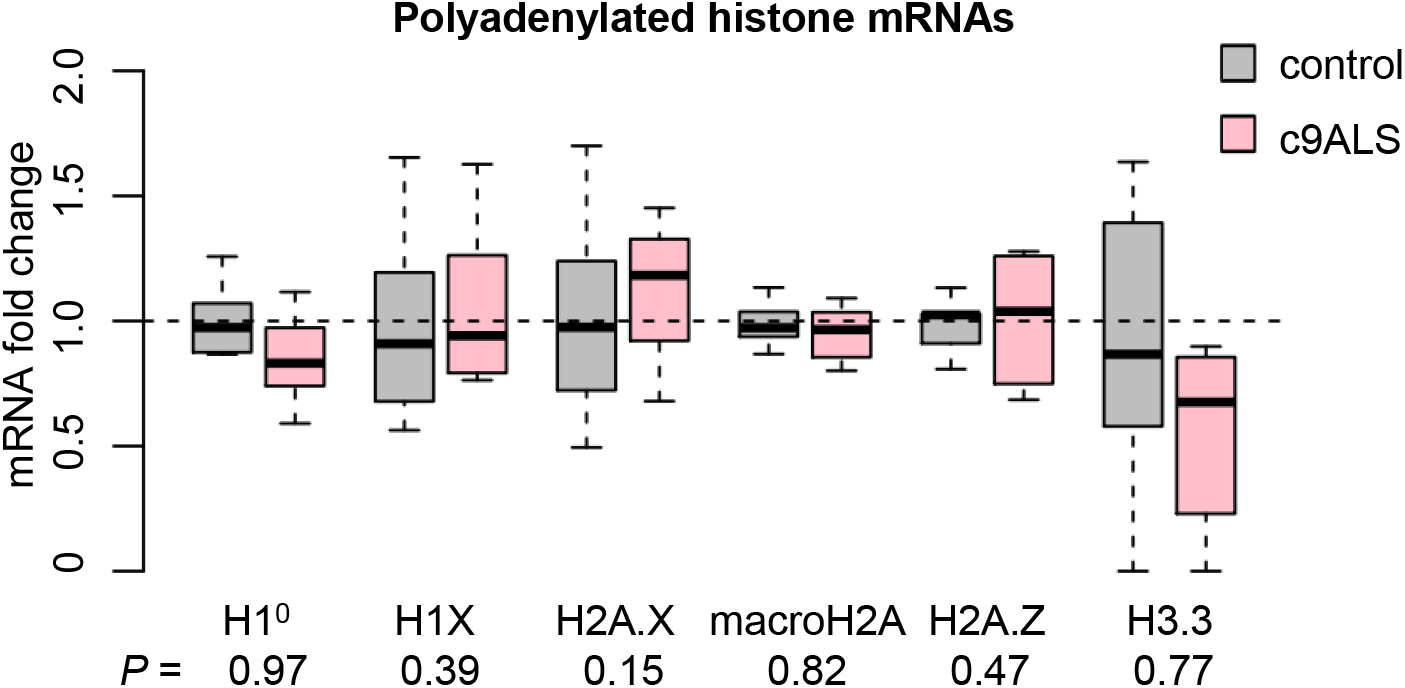
Expression of polyadenylated histone variants in c9ALS brains. Changes in mRNA abundance of noncanonical, polyadenylated histone mRNAs between c9ALS and controls. Expression levels were normalized to the means of control subjects. *P* values, one-sided Student’s t-tests.

**Figure 3 – figure supplement 1.**
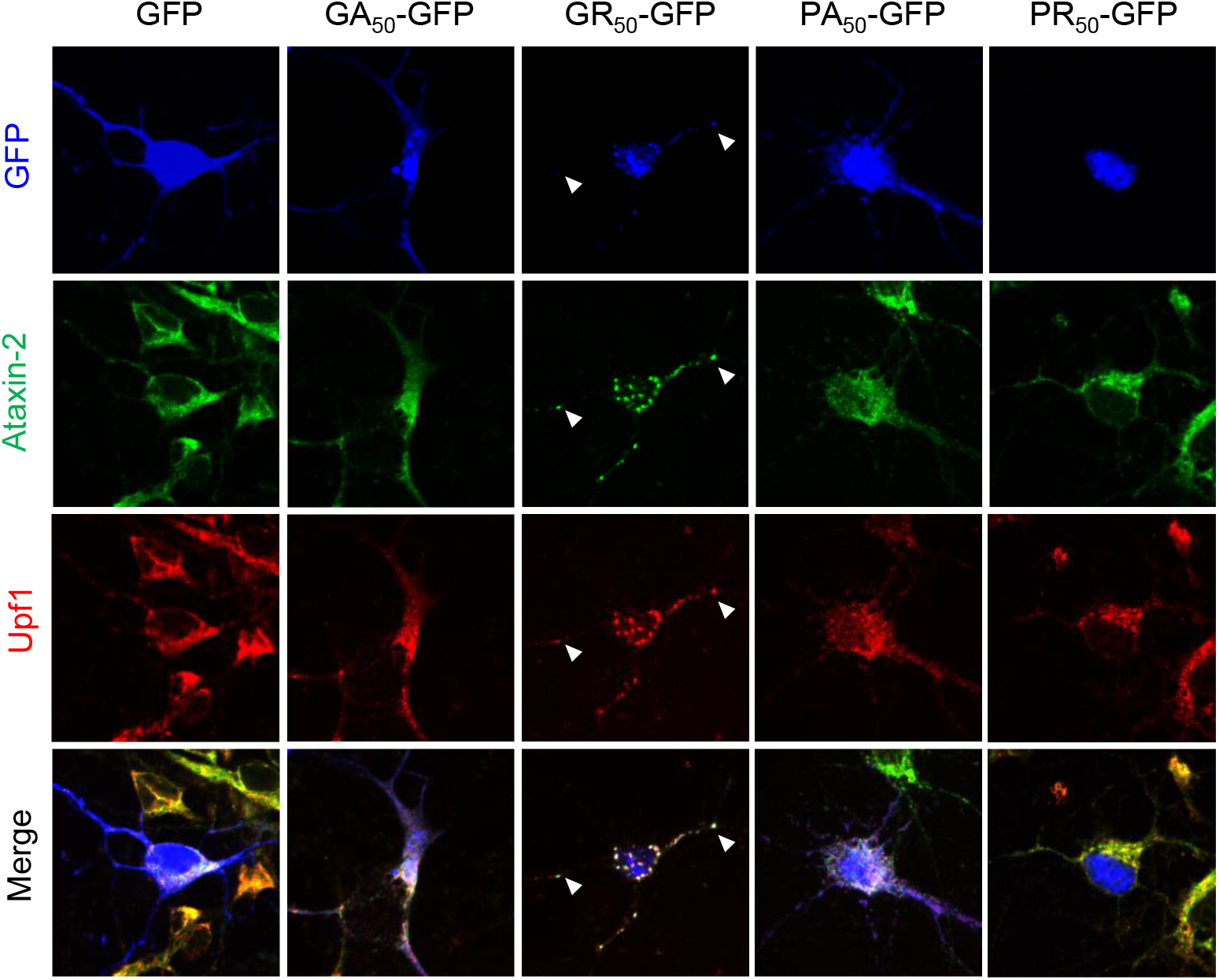
Poly(GR)-induces stress granules in primary neurons. Localization of Upf1 and Ataxin-2 in mouse primary cortical neurons expressing GFP or each DPR. Upf1^+^ Ataxin-2^+^ stress granules are indicated by arrowheads.

**Figure 3 – figure supplement 2.**
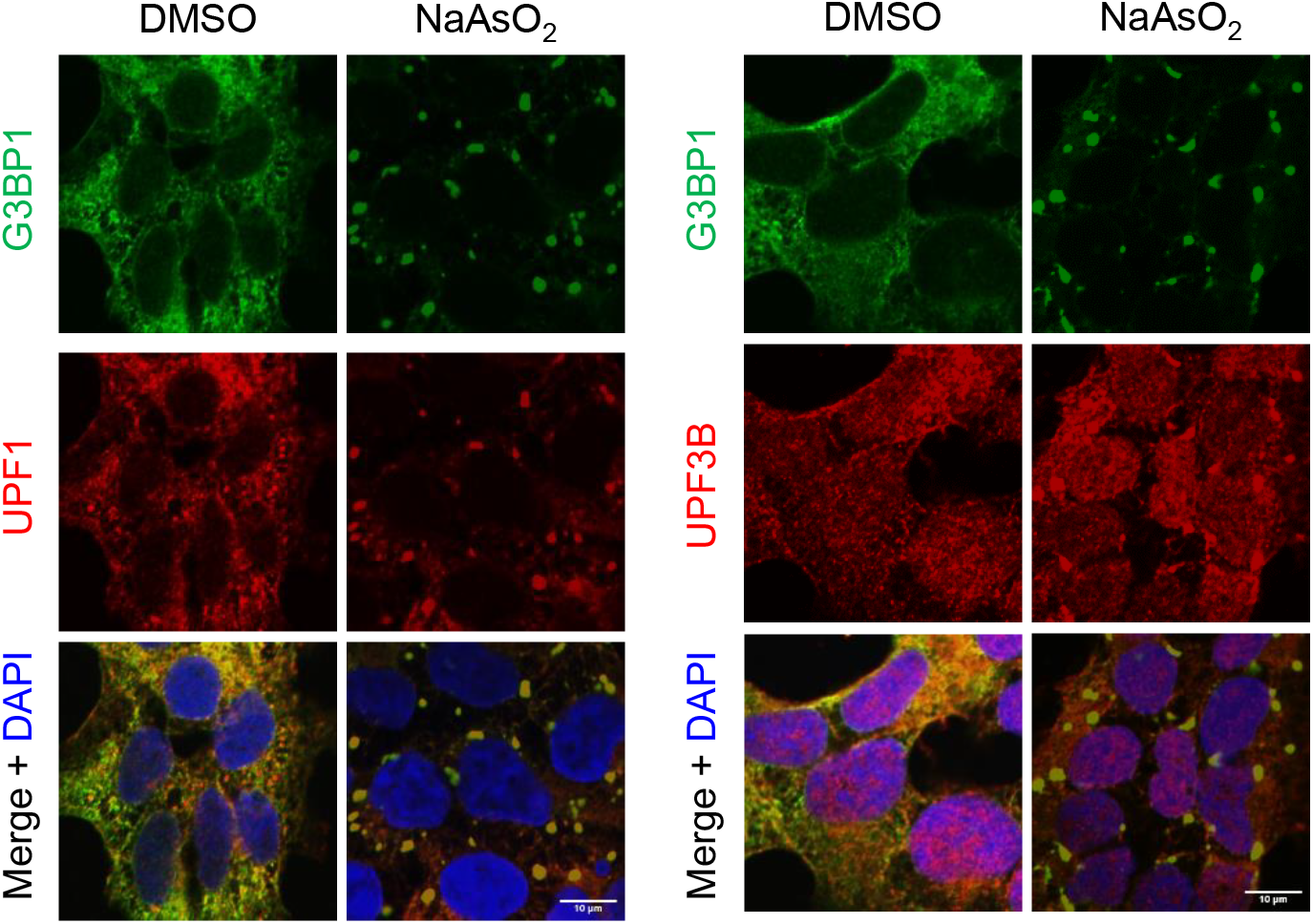
Recruitment of NMD factors to NaAsO_2_-induced stress granules. Localization of G3BP1, UPF1 (*left*) and UPF3B (*right*) in U-2 OS cells treated with DMSO or 0.5mM NaAsO_2_ for 30 min. Scale bars, 10 μm.

**Figure 3 – figure supplement 3.**
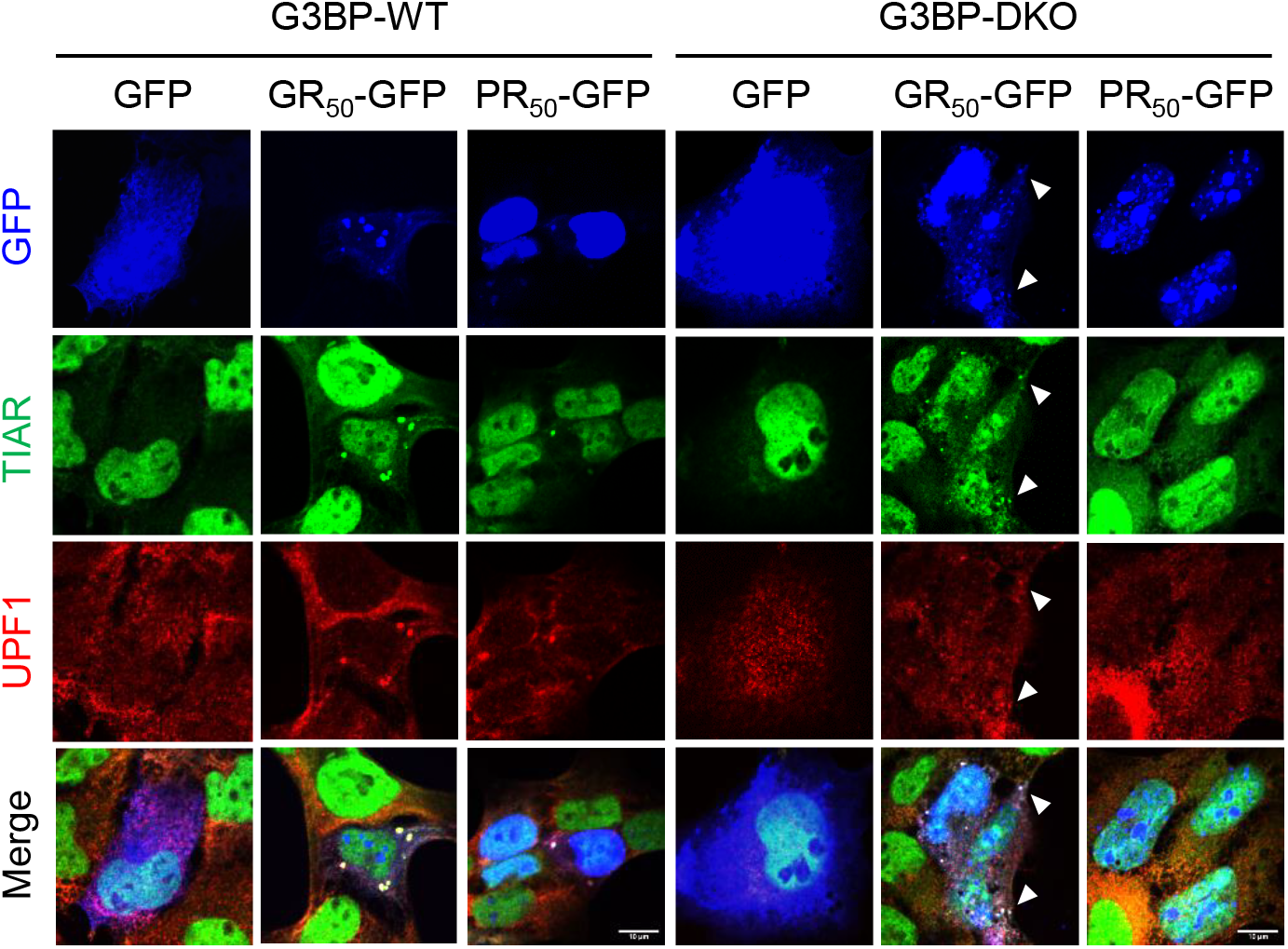
UPF1 localization in G3BP1/2-deficient U-2 OS cells. Localization of UPF1 and TIAR in G3BP-WT (*left*) and G3BP2-DKO (*right*) cells expressing either GFP, poly(GR), or poly(PR). Small TIAR^+^ UPF1^+^ granules in poly(GR)-expressing G3BP-DKO cells are indicated by arrowheads. Scale bars, 10 μm.

